# Implementing new portable touchscreen-setups to enhance cognitive research and enrich zoo-housed animals

**DOI:** 10.1101/316042

**Authors:** Vanessa Schmitt

**Affiliations:** Tiergarten Heidelberg gGmbH, Heidelberg, Germany; Centre for Organismal Studies (COS), University of Heidelberg, Heidelberg, Germany

**Keywords:** Touchscreen technology, Comparative Cognition, Animal-Computer-Interaction, Welfare, Primates, Birds, Enrichment, Social Group

## Abstract

To understand the evolutionary development of cognition, comparing the cognitive capacities of different animal species is essential. However, getting access to various species with sufficient sample sizes can be very challenging. Zoos, housing large ranges of animal taxa, would offer ideal research environments, but zoo-based studies on cognition are still rare. The use of touchscreen-computers to explore the cognitive abilities of nonhuman animals has shown to be highly applicable, and simultaneously offers new enrichment possibilities for captive animals. To facilitate zoo-based research, I here illustrate the assembly and usage of newly developed touchscreen-computer-systems (**Z**oo-based **A**nimal-**C**omputer-**I**nteraction System, **ZACI**), which can be used in various zoo environments and, importantly, with different taxa (e.g. primates, birds). The developed setups are portable, can be attached to various mesh sizes, and do not need any external power supply while being used. To evaluate the usability of the ZACI, they have been tested with experimentally naïve subjects of three great ape species (orang-utans, chimpanzees, gorillas) housed at Zoo Heidelberg, Germany, demonstrating to be easy to handle, animal-proof, and of great interest to the animals. Animals could be tested within their social group, as each subject had access to its own device during testing. To support the implementation of touchscreen-setups at other facilities, I also illustrate the training procedure and present first data on the apes’ performance in a simple object discrimination task. Portable touchscreen-setups offer the great possibility to enhance collaboration between zoos and researchers, allow a standardisation of methods, and improve data collection.

## Introduction

Comparing the cognitive abilities of different animal species to elucidate the evolutionary trajectories of cognitive development constitutes a promising research avenue (Herrmann et al. 2007; Schmitt et al. 2012; Benson-Amran et al. 2016;Vonk 2016; Whiten 2017), but often studies struggle with limited access to various species or subjects (MacLean et al. 2012; Tomasello & Call 2011; Thornton & Lukas 2012). Zoos, housing large ranges of animal taxa, would offer ideal research environments, but although cognitive research is increasingly taking place in zoos in recent years, especially in the US, it is still conducted at only a few facilities in Europe (Hopper 2017). Some zoos do not support basic scientific research, others do not have financial resources to hire scientific personal, and often logistical limitations prevent scientific research in zoos (see Hopper 2017 for a review).

Computerized technology provides well-established tools to conduct cognitive experiments with captive animals (e.g. Savage-Rumbaugh et al. 1986; Matsuzawa 1985; Bielick & Doering 1997; see alsoLeighty and Fragaszy 2003) and there is even a new research field emerging focussing on Animal-Computer-Interactions (ACI) (Pons et al. 2015; Mancini et al. 2017). In recent years, Lincoln Park Zoo in Chicago, USA, already established a comparative touchscreen research program with gorillas (*Gorilla gorilla gorilla*), chimpanzees (*Pan troglodytes*), and macaques (*Macaca fuscata*) (Egelkamp et al. 2016). In addition to primates, touchscreen experiments are now also being conducted with a large range of different animal species and taxa e.g. birds (*Nestor notabilis*, O’Hara et al. 2015), bears (*Ursus americanus,* Vonk & Beran 2012; *Helarctos malayanus*, Perdue 2016), dogs (*Canis lupus familiaries,* e.g. Zeagler et al. 2016), and even tortoises (*Chelonoidis carbonaria*, Mueller-Paul et al. 2014).

Touchscreen-technology offers a powerful research tool for various reasons. First, touchscreen-computers enable a valuable increase in accuracy and clarity of data collection, especially in comparison to manual tasks (e.g. Benson-Amram et al. 2016). Touchscreens facilitate very fine-grained measurements and accurate recordings of metrics like latency, improving cross-species comparisons (e.g. chimpanzees and humans, Inoue & Matsuzawa 2007). Second, data collection increases considerably, as it is possible to run hundreds of standardized and replicable trials in very short time spans (e.g. Fagot & Paleressompoulle 2009). Third, the variety of cognitive research questions, which can be explored, is nearly unlimited, and range from e.g. object, quantity or facial discrimination experiments (e.g. Vonk & Beran 2012; Micheletta et al. 2015; Johnson-Ulrich et al. 2016), examining the understanding of social interactions (e.g. Waller et al. 2016), evaluating emotional effects on cognition (e.g. Allritz et al. 2016), to virtually simulated reality experiments (Dolins et al. 2014; 2017). Fourth, when set up as an automated system, they do not need the constant interaction with a human experimenter, preventing unintentional biases such as cueing subjects’ responses, which, in turn, leads to more robust results (Clever Hans effect, e.g. Schmidjell et al. 2012). This little human interaction also facilitates the implementation of such test devices in zoological settings, where access to the animals and research personnel may be limited. When using multiple touchscreen-setups it is even possible to test animals individually, but while they are staying within their social group (Gazes et al. 2013; Fagot & Paleressompoulle 2009), or it can be tested how animals working in parallel influence each other (e.g. Martin et al. 2011; Schmitt et al. 2016).

Besides increasing the number of potential test subjects and improving data collection for cognitive research, conducting touchscreen studies at zoos may also contribute to enhance animal welfare (e.g. Perdue et al. 2012a). A recent study in zoo-housed crested macaques (*Macaca nigra*) already indicated that conducting scientific touchscreen studies positively influenced their wellbeing (Whitehouse et al. 2013), and so-called ‘cognitive enrichment’ is now being considered as an important factor in zoo welfare management (see Clark 2011 and 2017 for review). Interestingly, some zoos and sanctuaries already give their orang-utans access to Ipads as behavioural enrichment activity (Boostrom 2013, Wirman 2013).

Touchscreen-technology is thus increasingly applied to test the cognitive abilities of or enrich zoo-housed animals, but the actual setups used vary. Some studies use build-in systems (e.g. Ross 2009; Micheletta et al. 2015; Allritz et al. 2016), which often need expensive or laborious construction work that some zoos do not support. Furthermore, these build-in touchscreens are most of the time only accessible or used by a limited number of subjects or species per zoo (e.g. Perdue et al. 2012a; Micheletta et al. 2015). In contrast, portable or at least moveable touchscreen-systems can allow for a variety of species tested within the same zoo. Some studies place a single tablet or touch-monitor in front of the animals, but these setups often require close human contact, as e.g. holding the monitor in front of the subjects (e.g. Vonk 2003; Wirman 2013) or feeding the animals manually (e.g. Sonnweber et al. 2015; Altschul et al. 2017); procedures which some zoos do not approve. Therefore, at Zoo Atlanta, USA, researchers use stand-alone, portable touchscreen-setups that can be attached to the mesh of the animals’ enclosures and are equipped with food dispensers (Diamond et al. 2016; Perdue 2016). At Indianapolis Zoo, USA, C.F. Martin is also developing similar portable touchscreen-setups to work with great apes with improved technological components, as e.g. attaching a battery to allow a cordless usage (C.F. Martin, personal communication, June 2017). Detailed documentations on the construction of such touchscreen-setups are, however, not often published or only available for systems designed for laboratory purposes (e.g. Steurer et al. 2012, Calapai et al. 2016; see also http://lafayetteneuroscience.com/ for a primate-specific setup available for purchase), or describe build-in setups like the arena system (Martin et al. 2014), which are not easily implemented at most zoos.

To facilitate cognitive testing with zoo-housed animals, I here illustrate the assembly and usage of a newly developed portable touchscreen-setup, which can be used in various zoo environments (e.g. different mesh constructions) and, importantly, with different animal species (also non-primate). The setups are portable, can be attached to various mesh sizes, and do not need any external power supply while being used. To validate the usability of the touchscreen-setups, they have been tested with orang-utans (*Pongo abelii*), chimpanzees (*Pan troglodytes*), and gorillas (*Gorilla gorilla gorilla*) at Zoo Heidelberg (12 individuals, ages ranging from 5 to 45 years). The aim of this report is twofold. First, to illustrate the assembly of the setups to facilitate collaborations between researchers and zoos, and to enable a possible reproduction at other facilities. Second, to examine whether and how experimentally naïve subjects of three different great apes species interact with the new touchscreen-setups, and can be introduced to use such devices.

## Material & Methods

### The Zoo-based Animal-Computer-Interaction System (ZACI)

The development of the Zoo-based Animal-Computer-Interaction System (ZACI) was inspired by the portable touchscreen-setups used at Zoo Atlanta (personal observations and personal communication by R. Paxton Gazes, June 2014) and Indianapolis Zoo (C. F. Martin, personal communication, July 2014 and March 2015), but it integrates different technologies and newly developed design components, like the Electronic Control Unit (ECU, see below). The ZACI functions as a stand-alone, portable system, which can be attached to the outside iron bars of the animal’s enclosure. It is adjustable to different mesh sizes via adjusting joined hooks at the top and at the bottom of the apparatus **(Figure 1** and **supplemental information**). Furthermore, it does not need any external power supply while being used, reducing the risk of electricity cables within an animal’s proximity.

**Figure 1:**
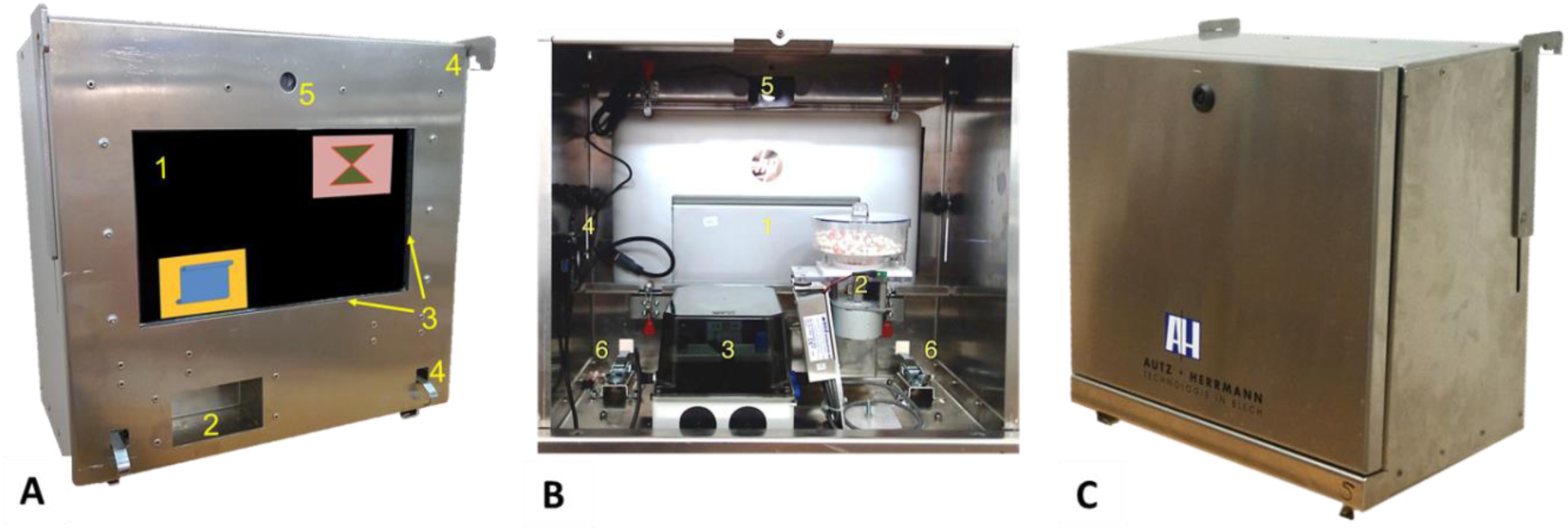
**A)** Front view of the touchscreen-system including a 15.6” laptop, which is located behind a 10mm Plexiglas^®^ panel (1), an opening for reward pellets (2), a 15.6” infrared touchframe surrounding the screen (3), hooks adjustable to different mesh sizes (4), and a webcam to film the subjects (5). **B)** Back view of the touchscreen-system showing laptop (1), food dispenser (2), the self-developed ECU with power supply (3), a USB-hub to connect the components (4), the webcam (5), and hooks (6). **C)** Back view of the setup with attached back cover.

The core of a unit is a 15.6” convertible laptop (HP ENVY x2 15-c000ng 2in1, by HP^®^ Inc., Palo Alto, CA) running Microsoft Windows^®^, which can be used as a tablet computer with a detachable keyboard (**Figure 1A & 1B**). However, any convertible laptop or tablet computer of respective size could be used to run the ZACI. The laptop can be slid in and out of the system to be recharged or programmed within seconds, using hooks attached to the aluminium casing. The detachable keyboard of the currently used laptop functions via Bluetooth and allows operating the laptop even when it is attached to the system. To protect the laptop from damage and dirt, a transparent Plexiglas^®^ panel (width 10 mm) is located directly in front of the screen and adjusted to the metal casing, i.e. the animals only touch at the Plexiglas panel and not the laptop. To register touches a 15.6” infrared (IR) touchframe (Model PPMT-IR-0156GR-WP, by KEYTEC^®^, Inc., Garland, TX) is surrounding the Plexiglas panel, facing the subjects, and is connected to the laptop via USB. The IR-touchframe technology allows the device to be used not only by touching it with a finger, but also by e.g. a bird’s beak, a tongue or a stick. In fact, anything penetrating the infrared barrier can be used to elicit a response of the touchscreen (Steurer et al. 2012). For rewarding correct trials, the unit contains a pellet dispenser (Model ENV-203-190IR, by Med Associates Inc., St. Albans, VT) ejecting standardized reward pellets (190 mg, manufactured by TestDiet^®^, St. Louis, MO) in different flavours (e.g. fruit punch, apple, banana, peanut butter) (**Figure 1B**). But other pellet dispensers and reward pellets, respectively, could be integrated into the system depending on the species being tested. The laptop and the food dispenser are coupled via a self-developed electronic control unit (ECU), containing the necessary electronics and a rechargeable battery (see **supplement**). Furthermore, each ZACI includes an additional webcam (Live! Cam Sync HD 720p, by Creative^®^, Dublin, Ireland), making it possible to film the animals while working on the screen. The laptop, IR-touchframe, ECU, and webcam are interconnected via an additional USB-hub (USB 3.1 Hub, by CSL-Computer GmbH & Co. KG, Hannover, Germany) with four interfaces attached to the inside of the metal casing.

All components of the ZACI are integrated in an aluminium casing (45cm x 50cm x 26cm, HxWxD) manufactured by a local company (Autz & Herrmann GmbH, Heidelberg, see **supplement** for more pictures on the assembly of the ZACI). Aluminium is much lighter than steel (a 100 x 100 x 5 mm board made of steel weighs 0.39kg, made of aluminium only 0.14kg), but offers the same stability and cleanliness. The ZACI equipped with all components weighs approximately 12kg, which is approximately half the weight of the Monkey CANTAB system sold by Lafayette Instrument^®^, and the portable touchscreen-setups used at Zoo Atlanta. An optional back cover protects the interior of the ZACI from liquids like water or urine and unwanted access **(Figure 1C**). The detailed construction plans of the aluminium casing are property of the company Autz & Herrmann GmbH, but further details and measurements can be made available on request. Furthermore, the company can easily produce the aluminium casing on request. All components of the ZACI have been manufactured to ensure that the tested individuals cannot dismantle the setups or hurt themselves. As the mesh of animal enclosures in European zoos often allows primate subjects to reach through with their arms, the metal casing is constructed as a smooth rectangular box, so the animals cannot hold onto and pull at parts of the setup when attached to the mesh. There are no loose parts in the proximity of the subjects. Furthermore, attaching the setup to the mesh can be performed with the arms of the experimenter located inside the metal casing (with the back cover removed), protecting the human from animals’ reach.

While the animals work at the units, the setups can be remotely controlled via Wi-Fi using an additional tablet computer (e.g. Microsoft Surface 3) to launch and control the experiments. The animal laptop and control tablet were coupled using the software TightVNC (http://www.tightvnc.com/). As Wi-Fi is not common in most animal holdings areas, we created a private Wi-Fi using a mobile router (Mobile WLAN router M7350 by TP-link Technologies Co., Ltd, Germany).

To program and run experimental procedures custom-made software was developed using the freely available programming language Java^TM^ (Oracle^®^). The software records the identity of the individual (manual input), the type of the stimuli used in the task, the area each stimuli was presented on at the screen, the reaction time of the subject (i.e. the latency to complete a trial), whether the response was correct or not, the exact point (with x and y coordinates) where the animal touched the screen, etc. The settings of each experiment could be easily adjusted via small property files (see https://en.wikipedia.org/wiki/.properties). Java is very independent of the operating system and hardware. Thus, the present experiments can run on Microsoft Windows^®^, Linux^®^ and Apple macOS^®^. Furthermore, other behavioural software (e.g. E-Prime^®^) could be used to run the ZACI. The only part of the newly developed software, which is crucial to control the pellet dispenser, is the low-level component controlling the USB relay card included in the self-developed ECU. This software code could also be integrated into other behavioural software, and can be accessed from the **supplement**.

### Subjects

Five gorillas (*Gorilla gorilla gorilla*), four chimpanzees (*Pan troglodytes*), and three orang-utans (*Pongo abelii*), participated in the study **(Table 1)** and were tested between August 2015 and June 2016 (the gorillas resumed training in June 2017). All species were living within their respective social groups housed at Zoo Heidelberg, Germany, and had access to indoor and outdoor exhibits. None of the subjects had ever participated in any touchscreen experiments before. The animals were not food or water deprived for testing. All testing was non-invasive and subjects participated voluntarily. All experiments followed the Guidelines for the Treatment of Animals in Behavioural Research and Teaching published by the Association for the Study of Animal Behaviour (http://asab.nottingham.ac.uk/downloads/guidelines2006.pdf).

**Tabel 1:**
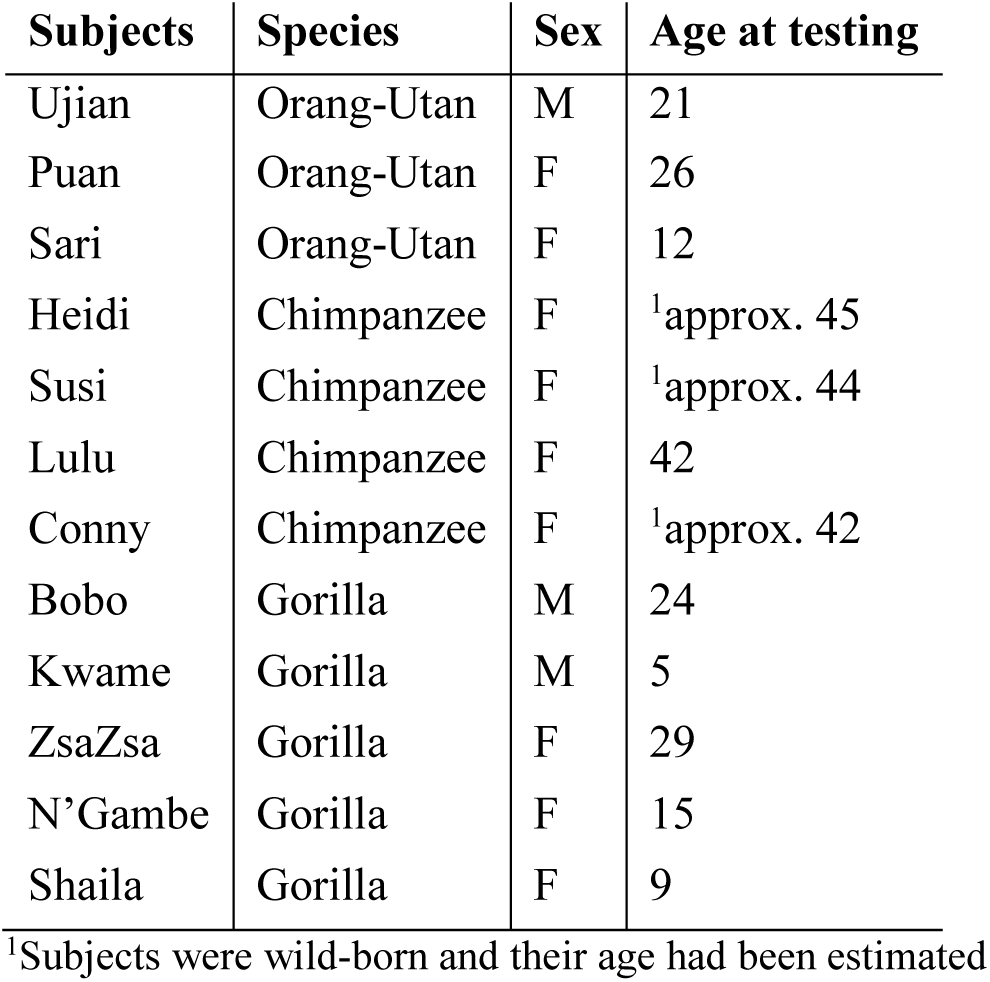
Name, species, sex, and age at testing of the subjects participating in the study.

### General Procedure

The orang-utans and chimpanzees were tested within their social groups and were not separated for the touchscreen tasks. The gorillas, however, were usually being separated in the indoor areas of their exhibit for afternoon feeding and cleaning, irrespective of the touchscreen testing (except for Kwame, the youngest subject, who always stayed with one of the females). Due to husbandry reasons, we began testing the gorillas during this time period, but after a couple of months they were also tested without being separated. As five units of the ZACI had already been built, it was possible to set up one unit for each subject. Each unit was attached to the outside of the iron bars of the animals’ indoor exhibits for approximately 45min per day, approximately 3 to 5 times a week depending on the least interference with regular husbandry procedures. At their first encounter of the touchscreens, we allowed the animals tested in groups to go to any unit. As soon as they had learned that touching the screens resulted in rewards, however, we trained the subjects that only one specific device would work for each animal. When subject A tried to activate subject B’s touchscreen, we unplugged the USB of the IR-screen. Thereby the animals learned within a couple of days which device they could work on. This procedure ensured reliable data collection in the following cognitive experiments.

### Experimental Procedure

#### Shaping

The initial experiment of the study trained the subjects to use the touchscreen and shaped their touching movements to reliably select small pictures on the screen. It consisted of six stages, guiding the animals from touching the whole screen to touching small pictures on the screen (**Figure 2A & Table 2**). In each Stage a random picture (clipart, geometrical forms or photograph) was presented at the screen. In Stage 1 touching any part of the screen, i.e. either the picture or the background area, resulted in a positive feedback: an immediate auditory feedback (“ping”, 865 msec), a green screen (1 sec), and a reward pellet. In all subsequent Stages, touching the black background area instead of a picture did not elicit any feedback. Instead, only touching the presented picture was rewarded. In Stage 6, a yellow square (representing a start signal) preceded each picture. The subjects had to touch the square and the following picture to receive a reward. In each Stage the position of the pictures on the screen was pseudorandomised (covering each part of the screen equally) and varied from trial to trial.

**Figure 2:**
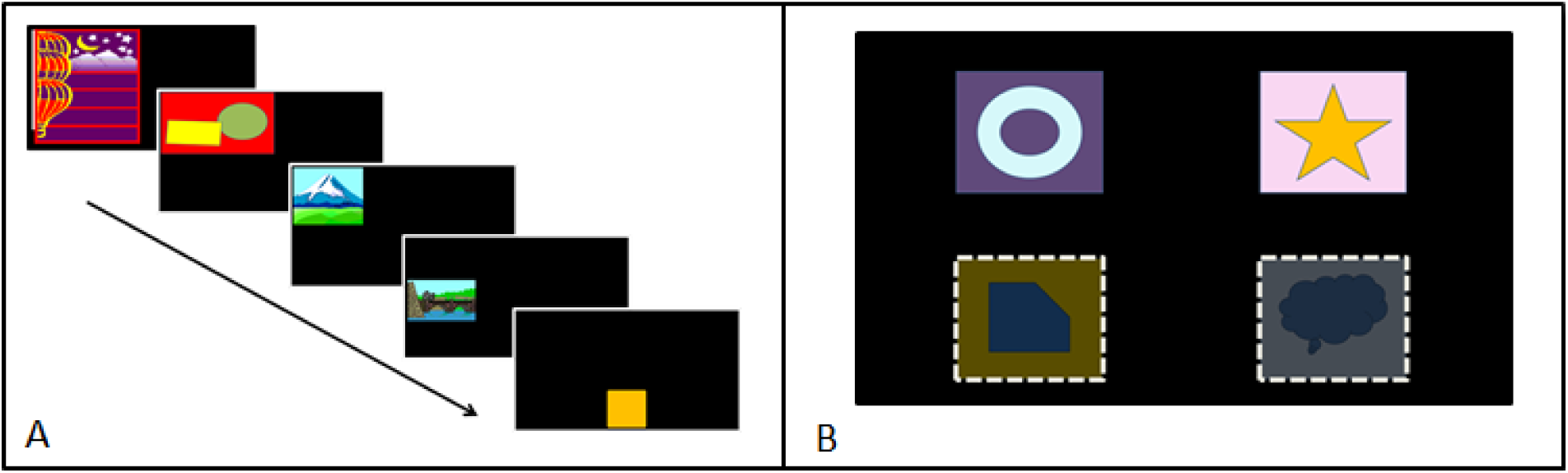
**A)** Representation of the successive stages of the *Shaping* experiment. The pictures the animals had to touch became smaller and ultimately preceded by a start signal (yellow square). **B)** Possible screen locations and picture samples of the *Object Discrimination* task. On each trial two pictures randomly appeared at the top, the bottom, only at the left or right side of the screen, or diagonally.

The experiment was designed to train each animal in a self-paced manner, so no predetermined number of trials in each stage was applied. Instead, each subject proceeded to the next stage after reaching a specific criterion, which demonstrated that the subject was able to focus on and work at the touchscreen for a certain amount of time (**Table 2**). For example, to pass Stage 1, the subject had to touch the screen for 12 consecutive trials needing not more than 6 seconds to complete each trial (the gorillas were allowed to take 8 seconds per trial as they needed more time to take the reward pellets out of the opening than the chimpanzees and orang-utans). In Stages 1 to 5 these reaction times were measured as the latency from the onset of each new picture on the screen until the subject touched the picture. From Stage 6 on, however, reaction time referred to the latency from touching the yellow start signal until touching the pursuing picture. In addition to their reaction time, the software also recorded the position of each touch at the screen, i.e. also at the unrewarded black background area, to examine the touching behaviour.

**Table 2:** Specifications for each Stage of the *Shaping* procedure, i.e. the area the subjects had to touch, the size of the presented pictures, and the criteria to pass each Stage.

| Stage | Required touch at | Picture Size (W x H; cm) | Criterion to pass each Stage |  |
| --- | --- | --- | --- | --- |
|  |  |  | *Number of Trials* | *Time per Trial* |
| 1 | Whole screen | 17 x 19.4 | 12 consecutive trials | 6 seconds (8 seconds for gorillas) |
| 2 | Picture | 17 x 19.4 | 20 consecutive trials | 6 seconds |
| 3 | Picture | 17 x 10 | 20 consecutive trials | 6 seconds |
| 4 | Picture | 10.5 x 10 | 20 consecutive trials | 6 seconds |
| 5 | Picture | 10.5 x 6 | 20 consecutive trials | 6 seconds |
| 6 | Start Signal + Picture | 10.5 x 6 | 20 consecutive trials | 6 seconds |
|  |  |  | + 10 trials | 3 seconds |

#### Object Discrimination

After the animals had successfully mastered the *Shaping* procedure they moved on to their first *Object Discrimination* task. In this experiment the subjects learned to discriminate between two pictures (S+ and S−). Touching S+ resulted in the already known auditory feedback (“ping”), a green screen (1sec), and a reward pellet. Touching S- resulted in a new negative auditory feedback (“buzz”, 300 msec), a black time-out screen (3sec) and no reward pellet. Each trial began with the start signal (yellow square). All subjects received the same picture pairs (i.e. random geometrical forms), but whether a picture was assigned correct (S+) or incorrect (S-) was pseudorandomised. The pictures where randomly presented at four possible screen locations to prevent the development of any side biases (**Figure 2B**). After reaching a success ratio of 90% within twelve trials, i.e. eleven out of twelve trials correct, a new pair of symbols appeared on the screen and the subject entered the next level. The probability to choose 11 out of 12 trials correct by chance is p = .003. The correct and wrong pictures did not share any specific features between the levels. I present the data for the first three picture pairs the subjects successfully discriminated.

## Results

### Usability of the ZACI

The newly developed portable touchscreen systems proved to be highly applicable for the use with great apes housed at Zoo Heidelberg. Even after several months of testing, the animals were not able to dismantle or destroy any parts of the setups. Each subject interacted with the new apparatus and most learned how to control the touchscreen through the mesh of their enclosures (**Figure 3**).

**Figure 3:**
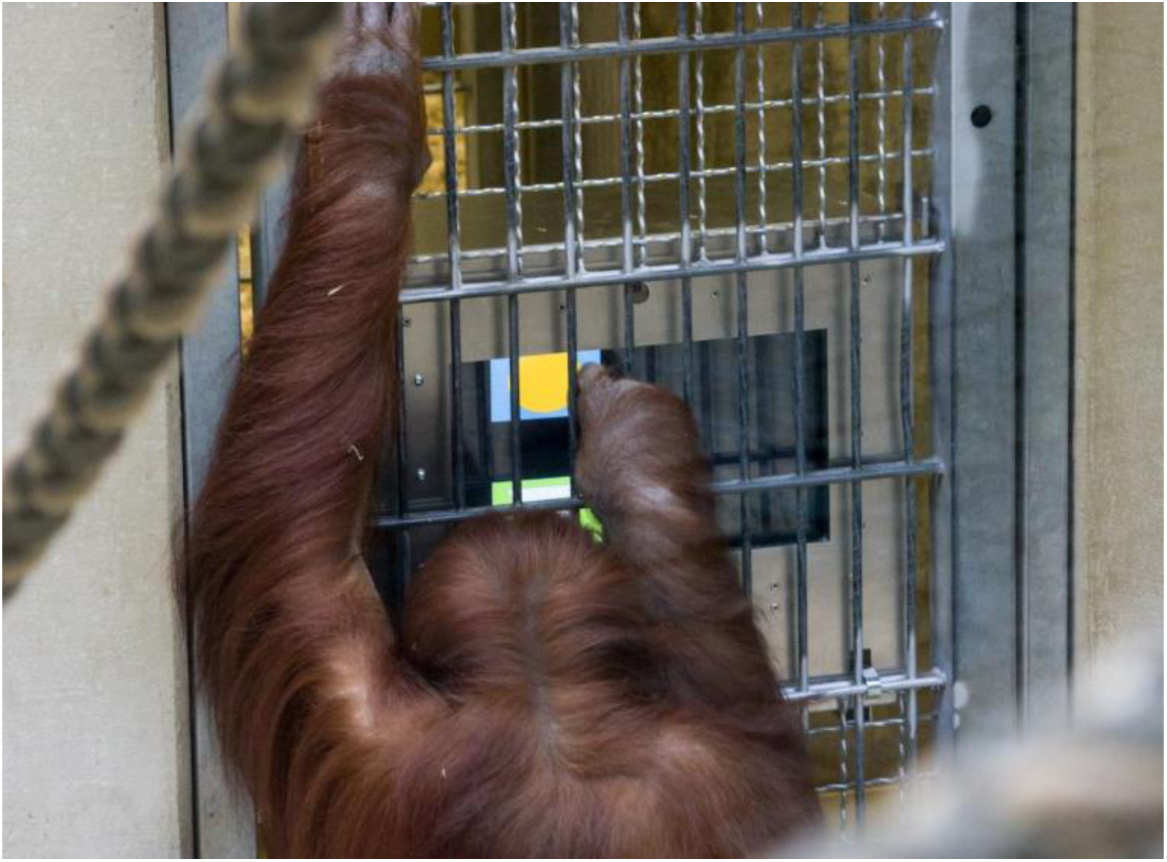
Orang-Utan Sari working at the Zoo-based Animal-Computer-Interaction system (ZACI). The system is attached to the outside bars of the animals’ enclosure and the subjects touch the screen through the mesh (Photo credit: Heidrun Knigge, Zoo Heidelberg).

### Shaping

After 2 months of training for each species, eleven of the twelve apes tested had passed the *Shaping* procedure. In fact, all orang-utans, all chimpanzees, and four gorillas had learned within 4 to 20 testing days to reliably touch small pictures on the screen (**Table 3**). The silverback gorilla still had to learn how to consistently touch small pictures, but all subjects showed large interest in the touchscreen systems and participated regularly in the experiments.

Initially, each subject should move to the next Stage of *Shaping* after reaching a specific criterion, i.e. passing a predetermined number of trials within a given timespan. However, during training some subjects had trouble meeting this criterion, as they either needed more time to take and eat the reward pellets or were generally more distracted. When a subject reliably touched the picture on the screen and only this timing issue interfered with advancement, difficulty was manually increased, i.e. picture sizes were decreased. This method enhanced efficiency of training for all subjects, except Bobo, the silverback gorilla. Manually increasing difficulty resulted in him stopping to participate. Only decreasing difficulty again, and then increasing it after 50 trials, got him back to work at the touchscreen. Notice, however, that manually increasing task difficulty was only applied until Stage 5. In Stage 6 each subject had to fulfil criterion, i.e. demonstrating a reaction time of less than 6 seconds for 20 consecutive trials and an additional 10 trials with less than 3 seconds per trial, for the *Shaping* procedure to be counted as passed.

**Table 3:**
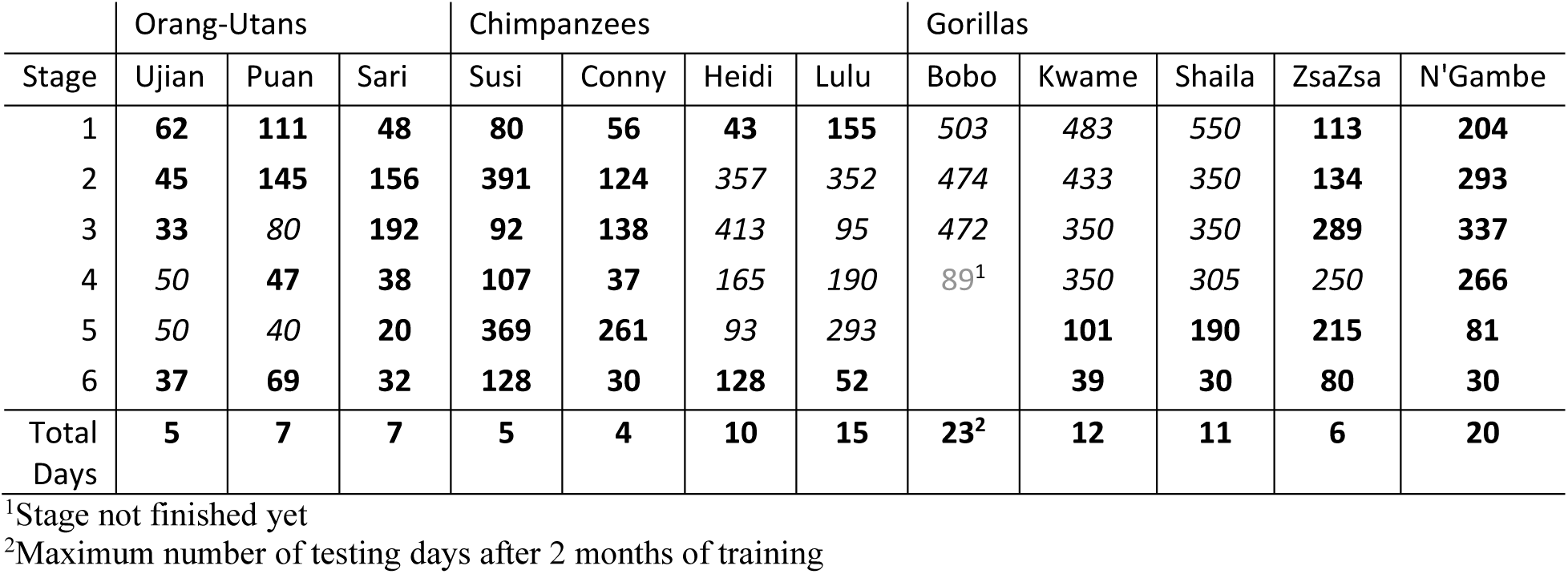
Number of trials and total number of testing days each subject needed to pass the six stages of *Shaping.* Numbers written in italics mean that the difficulty was manually increased to facilitate training.

|  | Orang-Utans |  |  | Chimpanzees |  |  |  | Gorillas |  |  |  |  |
| --- | --- | --- | --- | --- | --- | --- | --- | --- | --- | --- | --- | --- |
| Stage | Ujian | Puan | Sari | Susi | Conny | Heidi | Lulu | Bobo | Kwame | Shaila | ZsaZsa | N'Gambe |
| 1 | **62** | **111** | **48** | **80** | **56** | **43** | **155** | *503* | *483* | *550* | **113** | **204** |
| 2 | **45** | **145** | **156** | **391** | **124** | *357* | *352* | *474* | *433* | *350* | **134** | **293** |
| 3 | **33** | *80* | **192** | **92** | **138** | *413* | *95* | *472* | *350* | *350* | **289** | **337** |
| 4 | *50* | **47** | **38** | **107** | **37** | *165* | *190* | *89*<sup>1</sup> | *350* | *305* | *250* | **266** |
| 5 | *50* | *40* | **20** | **369** | **261** | *93* | *293* |  | **101** | **190** | **215** | **81** |
| 6 | **37** | **69** | **32** | **128** | **30** | **128** | **52** |  | **39** | **30** | **80** | **30** |
| Total Days | **5** | **7** | **7** | **5** | **4** | **10** | **15** | **23**<sup>2</sup> | **12** | **11** | **6** | **20** |

<sup>1</sup>Stage not finished yet
<sup>2</sup>Maximum number of testing days after 2 months of training

The computer also recorded where the animals touched the screen, i.e. whether and where they touched the unrewarded black background area. Although touching the background had no consequences for the subjects, it gave valuable information about any side preferences of the subjects’ touches. Sari’s heat map showed, for example, that she had difficulties to touch the part of the screen below the horizontal bar of the enclosure (see **Figure 4a**). This behaviour explained the large number of trials she needed to pass Stage 3, where the pictures only covered a quarter of the screen. After putting some honey on the lower part of the touchscreen, she quickly expanded her touches and finished Stage 4 to 6 needing the least number of trials of all subjects. In general, however, the bars in front of the touchscreen caused no difficulties for the subjects. All other subjects quickly learned to touch all parts of the screen (see **Figure 4b-d** for further examples).

**Figure 4:**
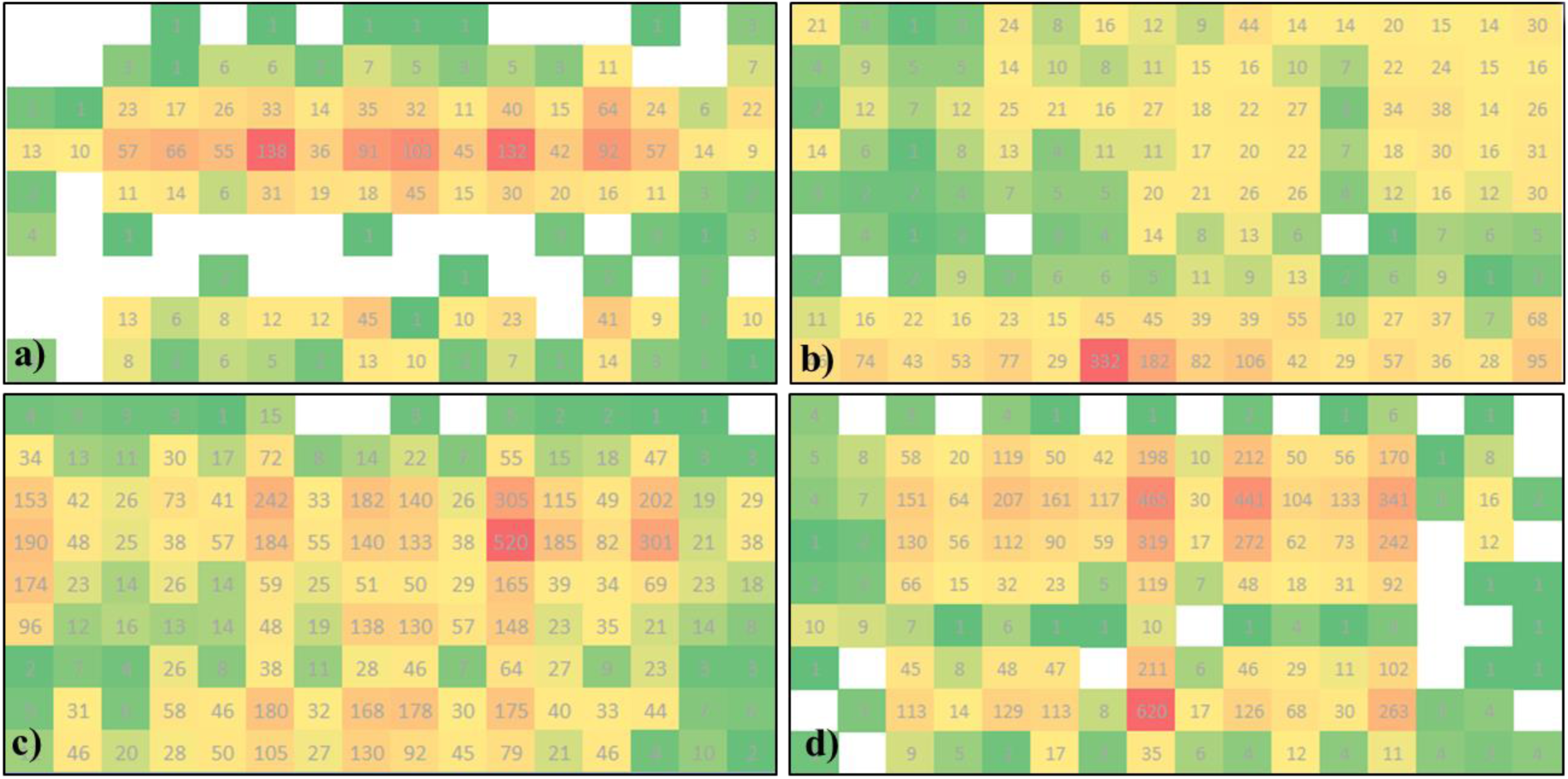
Distribution of touches on the screen during the *Shaping* task. The colour code indicates increased number of touches with increasing redness (heat map). The grey numbers give the total amount of touches at each square based on a 16 x 9 grid. **a)** Orang-Utan Sari, **b)** Orang-Utan Puan, **c)** Gorilla Kwame, **d)** Chimpanzee Conny.

### Object Discrimination

After the subjects had successfully passed the *Shaping* procedure they proceeded to their first *Object Discrimination* experiment, i.e. three orang-utans, four chimpanzees, and three gorillas participated (the gorilla N’Gambe had not finished *Shaping* yet). In this experimental paradigm, all subjects experienced a negative feedback, i.e. a buzz sound, no reward and a time-out, for the first time. However, all subjects continued to participate in the experiments and successfully discriminated three random picture pairs. Although visual inspection of **Figure 5** suggests that orang-utans performed slightly better than chimpanzees and gorillas, the results showed no significant species differences (two-way repeated measure ANOVA with species and Picture Pair as fixed factors and number of total trials as dependent variable, p = 0.209), but a significant effect of Picture Pair (p = 0.030), with subjects performing significantly better with Picture Pair 2 compared to Picture Pair 1 (p = 0.027). Instead, large individual differences have been observed, e.g. gorilla Kwame performing equally well as orang-utan Puan in discriminating picture pairs 2 and 3 (**Table 4**). As four possible screen locations were used, the development of any side biases was not observed.

**Table 4:** Number of wrong, correct, and total amount of trials each subject needed to pass the three *Object Discrimination* tasks.

| Species | Subject | Picture Pair | wrong | correct | total trials |
| --- | --- | --- | --- | --- | --- |
| Orang-Utan | Ujian | 1 | 46 | 63 | 109 |
|  |  | 2 | 63 | 53 | 116 |
|  |  | 3 | 24 | 32 | 56 |
|  | Puan | 1 | 62 | 108 | 170 |
|  |  | 2 | 7 | 22 | 29 |
|  |  | 3 | 7 | 29 | 36 |
|  | Sari | 1 | 107 | 150 | 257 |
|  |  | 2 | 132 | 139 | 271 |
|  |  | 3 | 66 | 85 | 151 |
| Chimpanzee | Conny | 1 | 289 | 338 | 627 |
|  |  | 2 | 204 | 216 | 420 |
|  |  | 3 | 453 | 563 | 1016 |
|  | Susi | 1 | 539 | 546 | 1085 |
|  |  | 2 | 185 | 108 | 293 |
|  |  | 3 | 191 | 202 | 393 |
|  | Heidi | 1 | 301 | 296 | 597 |
|  |  | 2 | 220 | 269 | 489 |
|  |  | 3 | 36 | 47 | 83 |
|  | Lulu | 1 | 126 | 161 | 287 |
|  |  | 2 | 134 | 124 | 258 |
|  |  | 3 | 241 | 224 | 465 |
| Gorilla | ZsaZsa | 1 | 689 | 794 | 1483 |
|  |  | 2 | 157 | 193 | 350 |
|  |  | 3 | 530 | 464 | 994 |
|  | Kwame | 1 | 265 | 293 | 558 |
|  |  | 2 | 14 | 36 | 50 |
|  |  | 3 | 24 | 36 | 60 |
|  | Shaila | 1 | 173 | 181 | 354 |
|  |  | 2 | 37 | 49 | 86 |
|  |  | 3 | 49 | 83 | 132 |

**Figure 5:**
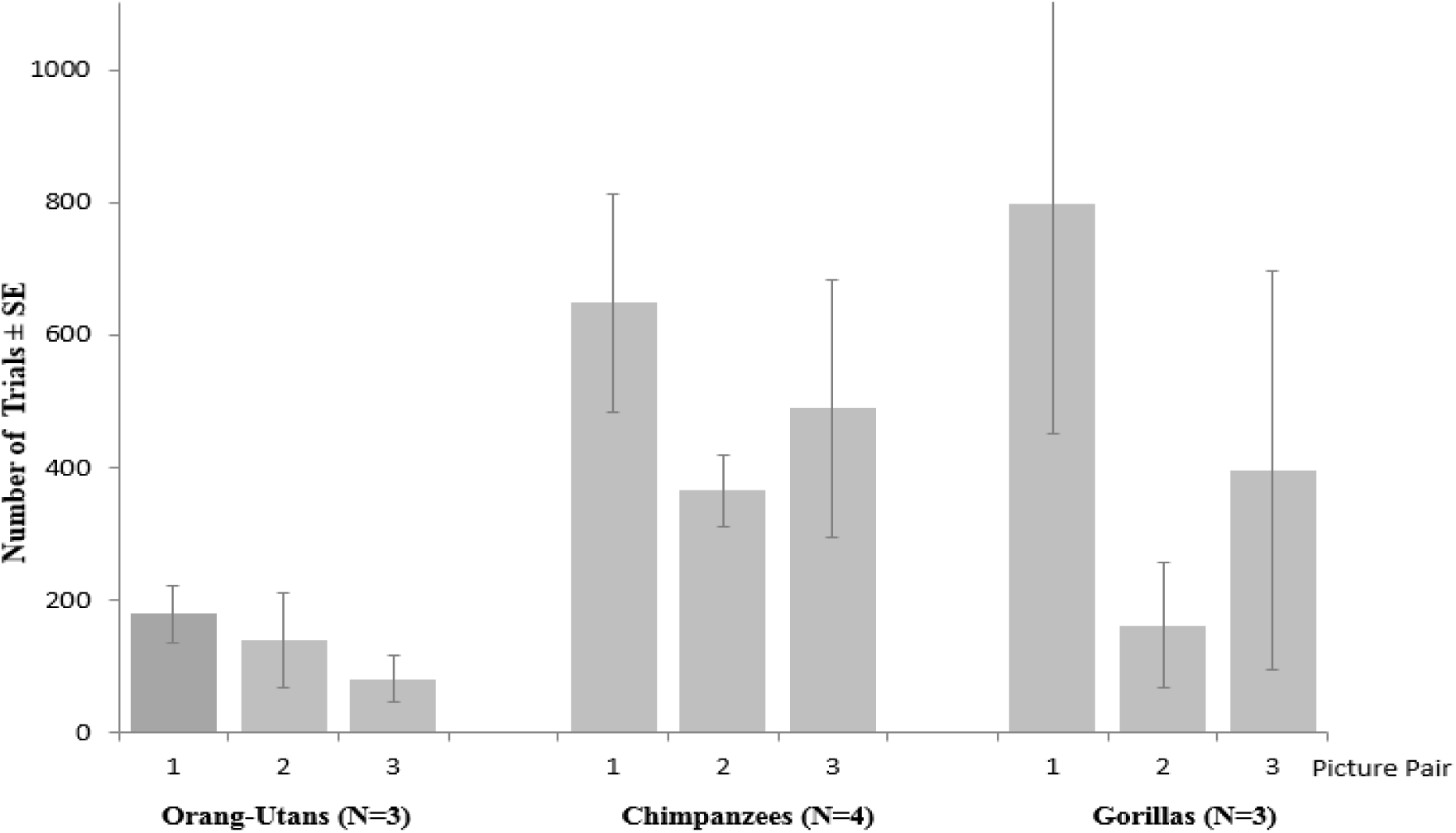
Mean number of trials ± standard error (SE) the apes needed to discriminate the first three random picture pairs in the *Object Discrimination* experiment.

## Discussion

The newly developed Zoo-based Animal-Computer-Interaction System (ZACI) proved to be highly applicable for work with zoo-housed primates. All subjects interacted with the touchscreen-systems on a regular basis and eleven out of twelve great apes learned to reliably use the setups within 4 to 20 days of testing. The silverback gorilla also showed large interest in the setups, but needs some additional testing days to finish the Shaping procedure. Using the whole area of the touchscreen through the mesh of their enclosure caused only some difficulties for one subject, but could easily be solved by putting honey at the screen. Placing the touchscreen on the outside of the mesh is also done at e.g. Zoo Atlanta or Marwell Zoo (Diamond et al. 2016; Perdue 2016; Micheletta et al. 2015), demonstrating a satisfactory touchscreen usability. Furthermore, even after several months of testing the apes were not able to dismantle or destroy any of the setups. Given the strength and intelligence of great apes, this confirms that the developed setups should also be safe to use with other animal species. The usage of an infrared touchframe allows the setup to be used also with non-primate species. A variety of studies already demonstrated that an infrared technology allows touchscreen usage by various animal species and taxa as for example birds, bears, dogs, and tortoises (e.g. Steurer et al. 2012; Vonk & Beran 2012; Mueller-Paul et al. 2014; Perdue 2016; Zeagler et al. 2016). The ZACI could therefore easily be introduced to a variety of animal species, also outside of a zoo setting. As the system can be closed with an optional back cover, protecting the interior from dirt and spray water, and does not need any power supply while being used, it could furthermore also be used outdoors, enabling an application at for example sanctuaries (as e.g. Wirman 2013). The training method applied in the *Shaping* task showed to be adequate for most subjects and elicited interest to continue touching the screen. Some subjects were not able to reach the predetermined temporal criterion in some Stages due to their slower touching frequencies, but manually increasing task difficulty considerably facilitated training. Only Bobo, the silverback gorilla, stopped working after manually increasing difficulty, and only participated again after resetting the Stage. Therefore, a slightly modified version of the training procedure might be more appropriate. Niessing and colleagues (2015) developed an automated training algorithm that uses a staircase procedure to train rhesus macaques (*Macaca mulatta*) the required touch at the screen. Their software either increases or decreases task difficulty based on the subject’s performance within the last 50 trials. Implementing a similar procedure might help to train animals without any human interference necessary and maybe even faster than has been accomplished in this study.

The quick acquisition of reliable touches by the animals now enables the collection of numerous data on their cognitive capacities. The first *Object Discrimination* experiment already demonstrates the usability of the setups to conduct experiments on comparative cognition. All subjects successfully discriminated three random picture pairs within only seven days of testing. The possible problem of developing side biases (see e.g. Allritz et al. 2016) could be solved by presenting the pictures not only at two, but at four possible locations, with the pictures also appearing either at only the left or right side of the screen, or diagonally. Interestingly, in the here presented *Object Discrimination* experiment the orang-utans performed slightly better than chimpanzees and gorillas. To declare any true species differences, however, larger sample sizes would be desirable. Establishing portable touchscreen-systems at other zoos with a greater collaboration between researchers could thus enhance data collection and enable robust analyses, helping to significantly advance the field of comparative cognition (see also Thornton & Lukas 2011).

In addition to increasing the number of potential test subjects for cognitive research, the usage of digital technology can serve as enrichment and even enhance the welfare of captive animals (Yamanashi & Hayashi 2011; Fagot et al. 2014; Bennett et al. 2016). The experimentally naïve apes of Zoo Heidelberg all interacted with the setups on a regular basis and some subjects even worked at the ZACI for 45min in each session, demonstrating the subjects’ large interest in the touchscreens and their enriching value. A new project at Zoo Melbourne even created digital projections for their orang-utans, allowing the apes and visitors to play interactive games (Webber et al. 2017). As a positive side effect, studies have shown that zoo visitors engaging with such technology show greater knowledge and interest in conservation issues, which could ultimately help to protect endangered species (e.g. Perdue et al. 2012b).

In conclusion, portable touchscreen setups offer the great possibility to significantly enhance data collection for scientific research on zoo-housed animals. In addition, they can improve animal welfare as they provide valuable enrichment to captive animals.

## Acknowledgments

Special thanks go to Christian Deutsch for developing the software used to program the experimental procedures and to integrate the technical components of the ZACI, and for helping to develop the electronic control unit. Many thanks also to Regina Paxton Gazes and Christopher Flynn Martin for sharing their knowledge on portable touchscreen devices and many helpful discussions during the construction. I am also grateful to student researchers, Marith Booijen, Jamie Dau, and Sarah Schilling, for their assistance with data collection. I thank the administration and employees of Zoo Heidelberg, especially Dr. Klaus Wünnemann and Sandra Reichler-Danielowski for supporting the study and providing the necessary infrastructure. Many thanks also to the animal caretakers of Zoo Heidelberg for their help in working with the great apes.

